# Human papillomavirus 16 E2 interaction with TopBP1 is required for E2 and viral genome stability during the viral life cycle

**DOI:** 10.1101/2023.01.11.523702

**Authors:** Apurva T. Prabhakar, Claire D. James, Christian T. Fontan, Raymonde Otoa, Xu Wang, Molly L. Bristol, Ronald D. Hill, Aanchal Dubey, Iain M. Morgan

## Abstract

CK2 phosphorylation of HPV16 E2 at serine 23 promotes interaction with TopBP1, and this interaction is important for E2 plasmid segregation function. Here we demonstrate that the E2-TopBP1 interaction is critical for E2 and viral genome stability during the viral life cycle. Introduction of the S23A mutation into the HPV16 genome results in a loss of E2 expression and viral genome integration during organotypic rafting. Co-culture of N/Tert-1+E2-S23A cells with J2 fibroblasts results in E2-S23A degradation via the proteasome, wild-type E2 is not degraded. TopBP1 siRNA treatment of N/Tert-1+E2-WT cells results in E2 degradation only in the presence of J2 cells demonstrating the critical role for TopBP1 in maintaining E2 stability. The CK2 inhibitor CX4945 promotes E2-WT degradation in the presence of fibroblasts as it disrupts E2-TopBP1 interaction. siRNA targeting SIRT1 rescues E2-S23A stability in N/Tert-1 cells treated with J2 fibroblasts, with an increased E2-S23A acetylation. The results demonstrate that the E2-TopBP1 interaction is critical during the viral life cycle as it prevents fibroblast stimulated SIRT1 mediated deacetylation of E2 that promotes protein degradation. This means that the E2-TopBP1 complex maintains E2 and viral genome stability and that disruption of this complex can promote viral genome integration. Finally, we demonstrate that HPV11 E2 also interacts with TopBP1 and that this interaction is critical for HPV11 E2 stability in the presence of J2 cells. Treatment of N/Tert-1+11E2-WT cells with CX4945 results in 11E2 degradation. Therefore, CK2 inhibition is a therapeutic strategy for alleviating HPV11 diseases, including juvenile respiratory papillomatosis.

**Importance:** Human papillomaviruses are pathogens that cause a host of diseases ranging from benign warts to cancers. There are no therapeutics available for combating these diseases that directly target viral proteins or processes, therefore we must enhance our understanding of HPV life cycles to assist with identifying novel treatments. In this report, we demonstrate that HPV16 and HPV11 E2 protein expression is dependent upon TopBP1 interaction in keratinocytes interacting with fibroblasts, which recapitulate stromal interactions in culture. The degradation of 16E2 promotes HPV16 genome integration, therefore the E2-TopBP1 interaction is critical during the viral life cycle. We demonstrate that the CK2 inhibitor CX4945 disrupts HPV11 interaction with TopBP1 and destabilizes HPV11 E2 protein in the presence of J2 fibroblasts; we propose that CX4945 could alleviate HPV11 disease burden.

## Introduction

Human papillomaviruses are causative agents in around 5% of all cancers, with HPV16 being the most prevalent type (1). Viral particles egress from the upper layers of infected stratified epithelium and can go on to infect stem like human keratinocytes in the basal layer (2, 3). Following mitosis of the infected cell, viral DNA is located in the host nucleus where cellular factors bind to the long control region (LCR, a non-coding part of the viral genome that regulates viral transcription and replication) and activate transcription from the viral genome (4–9). This results in production of viral proteins that are required for the viral life cycle. E6 and E7 are the primary viral oncogenes that drive proliferation of the infected cell (10), while the E1 and E2 proteins are replication factors (11, 12). The viral genome is an ~8kbp double stranded DNA episome and is replicated by E1-E2 in association with host cellular factors (13–15), and there are three phases to viral replication. First, there is an establishment phase immediately following infection when the viral genome stabilizes at around 50 copies per cell. Second, there is a maintenance phase where the viral genome copy number is maintained at around 50 copies per cell in the differentiating epithelium. Finally, in the upper layers there is an amplification stage of the viral life cycle where genome copy number per cell increases before encapsulation of viral genomes by the L1/L2 proteins to form viral particles which egress from the upper layers of the epithelium.

The E2 protein has multiple roles during the HPV16 life cycle (11). The carboxyl-terminal domain of E2 has a homodimerization domain and homodimers bind to 12bp palindromic DNA sequences in the LCR. Three of the E2 target sites surround the viral origin of replication, and via a protein-protein interaction E2 recruits the viral helicase to the A/T rich origin of replication where E1 forms a dihexameric helicase complex that replicates viral DNA in association with host factors (16, 17). E2 also regulates transcription from the viral LCR and is able to activate or repress transcription depending upon the levels of E2 protein, and can also regulate host gene transcription (18–23). The final role for E2 during the viral life cycle is to mediate plasmid genome segregation during cell division (24). In this process, during mitosis the E2 protein binds simultaneously to the viral genome and host chromatin, insuring that viral genomes reside in daughter nuclei following completion of mitosis. Several cellular factors have been implicated in mediating E2 association with mitotic chromatin to facilitate viral genome segregation, including BRD4 and ChlR1 (25–33). Recently we demonstrated that, for HPV16 E2, interaction with the cellular protein TopBP1 is required for E2 plasmid segregation function (34, 35). CK2 phosphorylation of E2 on serine 23 is required for HPV16 E2 interaction with TopBP1 (35). We also demonstrated that there is a TopBP1-BRD4 complex, and we propose that HPV16 E2 interaction with this complex is required for the plasmid segregation function of E2 and disruption of interaction with either TopBP1 or BRD4 could disrupt plasmid segregation function (35).

During the viral life cycle, double strand DNA breaks can result in integration of viral genomes into that of the host (36). Integration of viral DNA results in a worse clinical outcome, perhaps due to integration activating expression of endogenous adjacent genes such as *c-myc* (37–42). Integration also represents the end of the viral life cycle for the viral genome, therefore it is critical that it maintains an episomal genome, and the roles of E2 in replication and segregation are critical for this maintenance. In a recent manuscript we reported that introduction of the E2-S23A mutation (disrupting the E2-TopBP1 interaction) into the entire HPV16 genome resulted in a failure to detect E2 in human foreskin kerationcytes immortalized with the mutant (HFK+HPV16-E2-S23A) in organotypic rafts using E2 antibody immunofluorescent staining (35). The HPV16-E2-S23A rafts also had increased koilocytes, and invasion of the keratinocytes into the collagen-fibroblast plug that act as a stroma during the rafting process (35). This more aggressive behavior could be indicative of viral genome integration due to a loss of E2 protein. We therefore investigated the role of the E2-TopBP1 interaction in maintaining E2 and viral genome stability during the viral life cycle.

Here we demonstrate that, during the HPV16 life cycle in organotypic raft cultures, HFK+HPV16-E2-S23A cells had no detectable E2 protein expression, and that the viral genome is mostly integrated. Using N/Tert-1 cells stably expressing E2-WT and E2-S23A we demonstrate that, in the absence of binding to TopBP1, the E2 protein is targeted for proteasomal degradation in the presence of mouse or human fibroblasts. This degradation is due to enhanced deacetylation of E2 by SIRT1 when E2 is not interacting with TopBP1. The CK2 inhibitor CX4945 disrupts the E2-TopBP1 interaction (35) and promotes E2 degradation in the presence of fibroblasts. The results demonstrate that the E2-TopBP1 interaction is critical for the HPV16 life cycle.

The TopBP1 interacting sequence on HPV16 E2 is highly conserved in HPV11 E2. Thus, we also demonstrate that there is an interaction between HPV11 E2 and TopBP1. siRNA TopBP1 knockdown or treatment with CX4945 results in partial degradation of the HPV11 E2 protein in the presence of J2 fibroblasts. We propose that CX4945 is a potential candidate for the treatment of HPV11 diseases such as juvenile respiratory papillomatosis. The reduction in E2 protein level would reduce HPV11 replication resulting in less viral production and alleviation of disease.

## Results

### HPV16 E2 interaction with TopBP1 is required for E2 protein and viral genome stability in human keratinocytes interacting with J2 fibroblasts

Previously we demonstrated that there was a lack of detectable E2 expression in HFK+HPV16-E2-S23A in organotypic rafts when compared with HFK+HPV16 (wild type) by immunostaining (35). The HFK+HPV16-E2-S23A raft also had a more transformed phenotype with increased koilocytes and invasion into the collagen-fibroblast “stromal” plug, suggesting that the lack of E2 expression had promoted viral genome integration (35). To test this, protein and DNA was extracted from organotypic rafts and the expression of E2 and the viral genome status investigated. Figure 1A demonstrates that in HFK+HPV16-E2-S23A rafts there is no detectable E2 expression. Lanes 1 and 2 are N/Tert-1+Vec (cells G418 selected following pcDNA transfection) and N/Tert-1+E2 (stably expressing E2) respectively. Lane 3 is HFK+HPV16-E2-S23A, and lane 4 HFK+HPV16 (E2-WT). There is detectable E2 expressed in N/Tert-1+E2 and HFK+HPV16, but not in HFK+HPV16-E2-S23A, agreeing with our previous immunostaining data. To confirm that the HFK+HPV16-E2-S23A cells were expressing viral proteins, we tested for E7 expression and could see no difference in E7 protein levels between HFK+HPV16 and HFK+HPV16-E2-S23A. To confirm that viral replication was not occurring in the HFK+HPV16-E2-S23A cells, we blotted for γH2AX, a marker of the DNA damage response (DDR) induced by HPV16 replication. There is detectable γH2AX in HFK+HPV16 cells (lane 4), but not in HFK+HPV16-E2-S23A.

**Figure 1.**
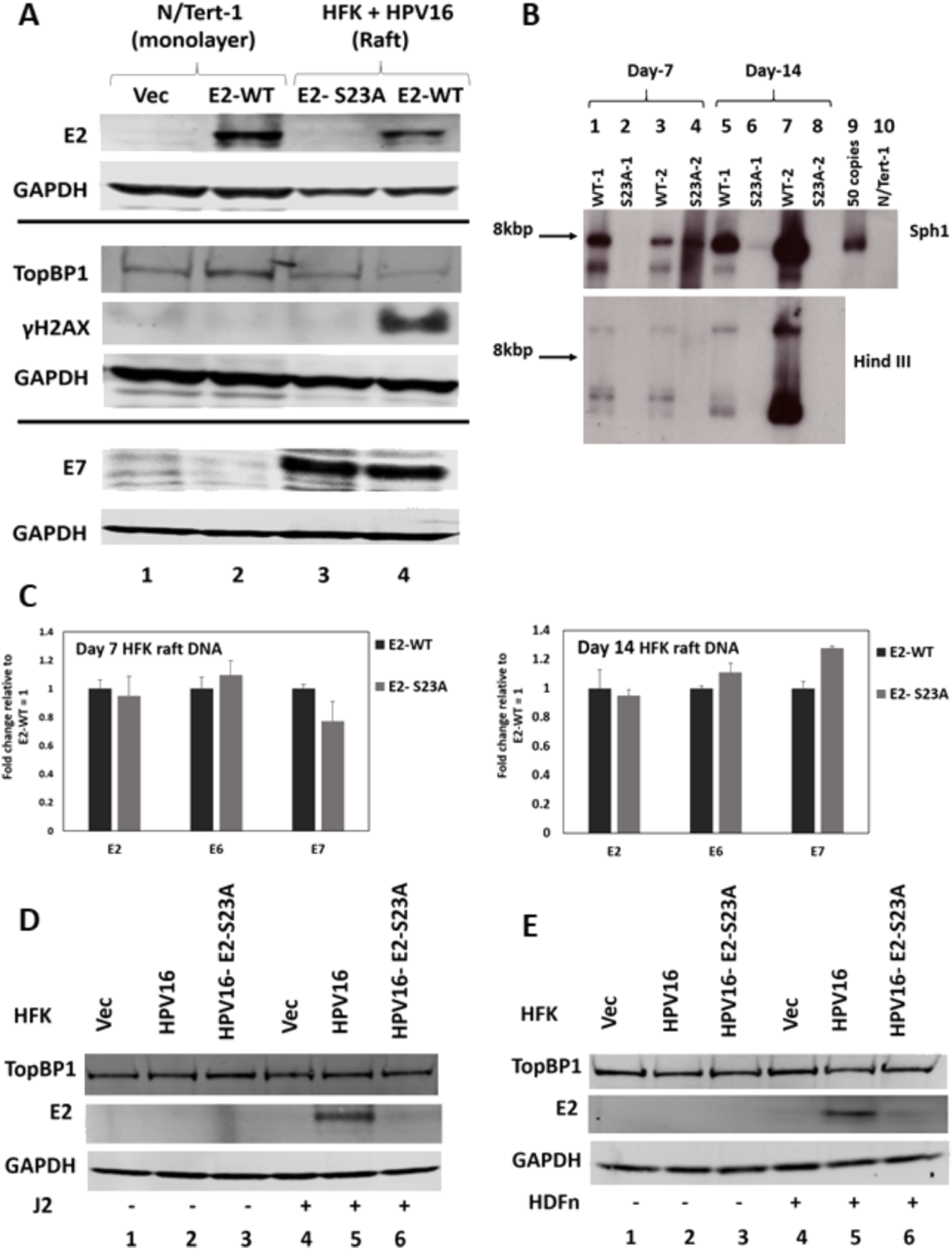
Interaction with TopBP1 is required for E2 protein stability and episomal viral genome maintenance during the viral life cycle. A. Protein extracts were prepared from the indicated cell lines and blotted with the antibodies listed on the left. Three western blots were carried out and the GAPDH control for each is bracketed with the antibodies used. B. DNA was harvested from HFK+HPV16 (WT) and HFK+HPV16-E2-S23A (S23A) organotypic raft cultures at day-7and day-14 after initiating of rafting. The DNA was digested with Sph1 (upper panel, cuts the HPV16 genome once) or HindIII (lower panel, does not cut the HPV16 genome) and Southern blotting carried out for the HPV16 genome. C. To confirm that the HFK+HPV16-E2-S23A DNA samples retained viral DNA, the indicated probes were used to monitor for the presence of HPV16 DNA using real-time PCR. Results were standardized to GAPDH and presented relative to HFK+HPV16 E2-WT equaling 1. D. N/Tert-1 Vec (control cells with no viral genomes but G418 selected), HFK+HPV16 (HPV16) and HFK+HPV16 E2-S23A (HPV16-E2-S23A) were cultured with and without J2 fibrobasts and protein extracts blotted with the indicated antibodies. E. The experiment in D. was repeated using neonatal human dermal fibroblasts.

The lack of viral replication markers, increased markers of a more transformed phenotype, and reduced E2 expression, suggested that the HPV16-E2-S23A genome had integrated. To investigate this, 7-day and 14-day raft cultures had DNA extracted. At 7-days full differentiation has not occurred, whereas 14-days gives time for full differentiation (as seen by production of a top layer of keratin) and viral genome amplification. Figure 1B details Southern blots from 2 independent HFK donors immortalized with HPV16 or HPV16-E2-S23A. In the upper panel the extracted DNA is digested by Sph1, a viral genome single cutter, and in the lower panel with HindIII, a viral genome non-cutter. With Sph1, in both WT-1 and WT-2 (HFK+HPV16) samples at 7-days (lanes 1 and 3 respectively) there is a detectable 8kbp band indicating that there are episomal genomes present. This is confirmed as at 14-days both WT-1 and WT-2 DNA levels are increased (lanes 5 and 7, respectively) due to viral genome amplification, which can only occur with episomal genomes. For S23A-1 there is no detectable 8kbp band at day-7 (lane 2) and a faint band at day-14 (lane 6) suggesting that the majority of the DNA has become integrated after 7-days and that a residual episomal sub-set of genomes is amplified between days 7 and 14, allowing detection of the faint band. This 8kbp band could also be due to the integration of HPV16 multimer genomes. For S23A-2 there is a different pattern. At day-7 there is a smear of DNA with a concentration around 8kbp, suggesting the beginning of integration (lane 4). At day-14 there is no detectable 8kbp band suggesting that most DNA has become integrated (lane 8). It should be stressed here that these HFK, prior to rafting, were grown independently of J2 expression and we have demonstrated that prior to rafting, there is episomal DNA in all of the HFK+HPV16 and HFK+HPV16-E2-S23A samples (35). In the uncut samples (lower panel), there is evidence of episomal viral genomes with the WT samples (three DNA bands are detected) and no evidence of episomal DNA with the S23A samples. Figure 1A showed that the HFK+HPV16-E2-S23A sample expresses equivalent levels of E7 protein when compared with HFK+HPV16, demonstrating that viral DNA must be present. To confirm this we carried out real-time PCR with E2, E6 and E7 primers (Figure 1C) and demonstrated the presence of viral DNA in WT-1 and S23A-1 samples. Results were normalized to GAPDH and expressed relative to E2 levels in the HFK+HPV16 WT cells. There was no statistically significant difference in the DNA levels between the WT-1 and S23A-1 samples.

We next grew one set of the HFK+HPV16 WT and E2-S23A cells in the presence and absence of mitomycin C treated J2 fibroblasts (Figure 1D). From now on, all fibroblasts used were treated with mitomycin C. Noticeably, in the absence of fibroblasts, there was poorly detectable levels of E2 protein expression, despite there being episomal viral genomes in the WT and E2-S23A lines (lanes 2 and 3 (35)). The addition of fibroblasts resulted in detection of E2 protein in WT cells (lane 5) but poor expression of E2-S23A (lane 6). These results suggest that the presence of fibroblasts changes the mode of HPV16 replication. In the absence of fibroblasts there are low levels of E2 protein and episomal genomes in WT and E2-S23A cells. In the presence of J2 cells there is an induction of E2 in WT cells and a lack of E2-S23A expression that results in episomal and integrated genomes, respectively. We therefore propose that culturing and study of HFK cells immortalized by HPV16 should always be carried out in the presence of fibroblasts. To confirm that the results with the J2 (mouse) fibroblasts were relevant to an actual HPV16 infection, we repeated the experiments with neonatal human dermal fibroblasts (HDFn) and obtained identical results (Figure 1E).

We have generated N/Tert-1 cells stably expressing E2-S23A (N/Tert-1+E2-S23A) (34, 35). Figure 2A demonstrates that, when J2 cells are added to N/Tert-1+E2-S23A, the E2 protein becomes undetectable by western blotting. There is no detectable E2 protein present in N/Tert-1+Vec (pcDNA control) cells with or without J2 cells (lanes 1 and 4). With N/Tert-1+E2 (wild type E2), E2 levels remains constant irrespective of J2 cells (lanes 2 and 5). However, in N/Tert-1+E2-S23A cells, the addition of J2 fibroblasts results in a reduction in detectable E2 levels (compare lanes 3 with 6). This was repeated an additional two times and quantitated (Figure 2B). TopBP1 levels are unchanged by either E2 expression or the addition of J2 cells. The differences in E2-S23A protein levels was not due to differences in E2 RNA levels following the addition of J2 cells (Figure 2C). To confirm that the addition of the fibroblasts was not regulating the differentiation of the N/Tert-1 cells, we determined that involucrin RNA levels were unchanged by the presence of J2 cells (Figure 2D). We next confirmed that HDFn generated identical results to mouse fibroblasts in these N/Tert-1 cells with regards regulation of E2 protein and RNA levels (Figures 2E, 2F and 2G).

**Figure 2.**
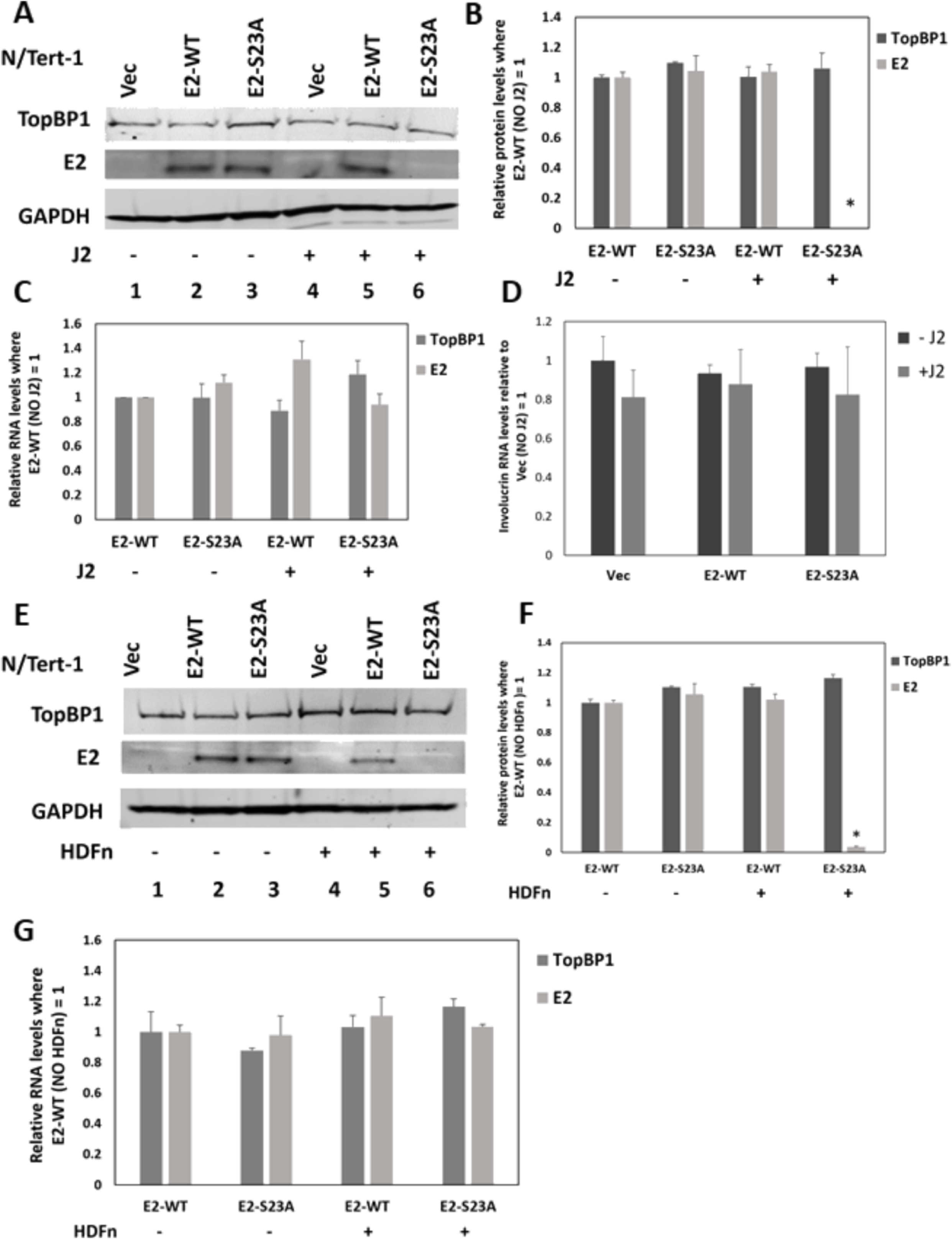
In N/tert-1 cells stably expressing E2-WT or E2-S23A, the addition of J2 fibroblasts selectively reduced E2-S23A protein levels. A. The indicated protein extracts were western blotted and probed with TopBP1, E2 or GAPDH. B. The experiment in A was repeated and the levels of E2 protein quantitated. * indicates a statistically significant reduction in E2-E23A protein levels in the presence of J2 cells when compared with no J2, p-value < 0.05. C. There was no significant change in E2 or TopBP1 RNA levels following the addition of J2 cells. D. The addition of J2 fibroblasts did not significantly change the expression of the differentiation marker involucrin. E. F. and G. are repeats of A. B. and C. using neonatal human dermal fibroblasts (HDFn) generating essentially identical results.

To confirm that the reduction in E2-S23A levels following addition of J2 cells was due to reduced interaction with TopBP1, we used siRNA against TopBP1 and demonstrated that reduction of TopBP1 reduced E2-WT levels in the presence of J2 in N/Tert-1 cells (Figure 3A). E2-WT levels are comparable between the J2 + and – samples (compare lane 2 with 4). In the absence of J2 cells, there was no dramatic difference in E2-WT levels when TopBP1 protein levels are reduced by siRNA targeting (compare lane 4 with lane 5). However, in the presence of J2 cells, reduction of TopBP1 levels resulted in a reduction in E2 protein levels (compare lane 2 with lane 3). For the E2-S23A mutant, siRNA targeting of TopBP1 did not change the protein levels in the absence of J2 cells (compare lane 8 with lane 9). The addition of J2 cells resulted in a reduction in E2-S23A protein levels irrespective of TopBP1 expression (lanes 6 and 7). This experiment was repeated and quantitated demonstrating a significant reduction in E2-WT protein levels in the presence of J2 cells when TopBP1 levels were reduced (Figure 3B). To confirm that this was not an off-target effect of the TopBP1 siRNA, we repeated the protein expression studies with a second TopBP1 siRNA and obtained identical results; a significant reduction in E2-WT protein levels in the presence of J2 cells and the absence of TopBP1 expression (Figures 3C and 3D). Removal of TopBP1 expression or addition of J2 fibroblasts did not alter the levels of E2 RNA (Figure 3E, results for TopBP1 siRNA A).

**Figure 3.**
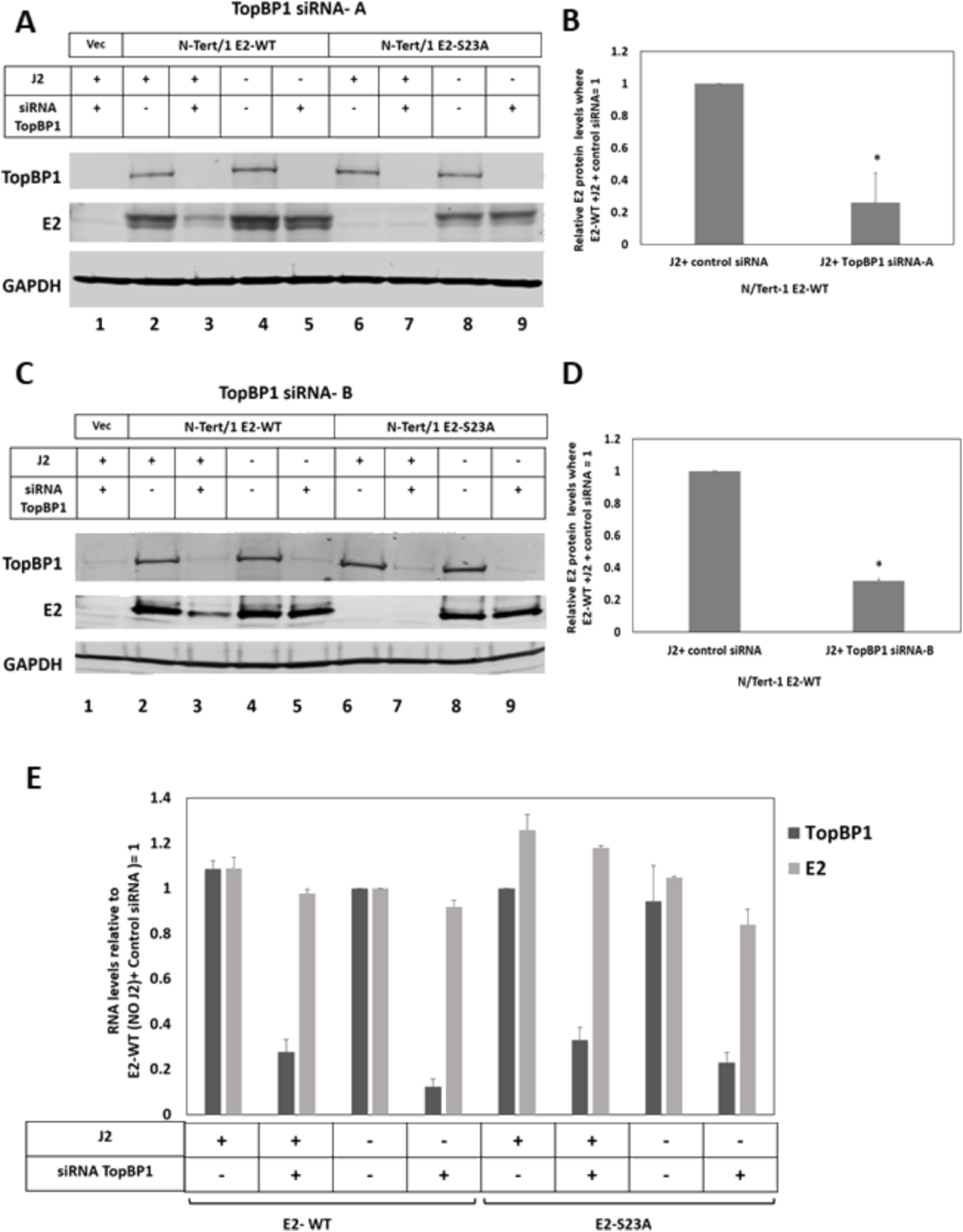
Knockdown of TopBP1 reduces E2-WT protein levels in the presence of J2 fibroblasts. A. The indicated cells were treated with scrambled siRNA or siRNA targeting TopBP1 in the presence or absence of J2 fibroblasts. Protein extracts were prepared and western blotting used to monitor TopBP1 and E2 protein levels, GAPDH was used as a loading control. B. The experiment shown in A was repeated and quantitated and the levels of E2-WT protein in the presence or absence of TopBP1 with J2 fibroblasts calculated. C-D. These figures represent a repetition of the TopBP1 siRNA experiments using an alternative TopBP1 targeting siRNA. In B and C, * indicates a statistically significant reduction in E2 protein levels when TopBP1 expression is knocked down in the presence of J2 fibroblasts, p-value < 0.05. E. Neither the TopBP1 siRNA treatment nor the addition of J2 fibroblasts statistically altered E2-WT or E2-S23A RNA expression levels. The results shown are a summary from two independent RNA preparations using TopBP1 siRNA A.

### Interaction with TopBP1 prevents E2 deacetylation by SIRT1 in the presence of J2 fibroblasts

Having established that interaction with TopBP1 is required for E2 stability in the presence of J2 fibroblasts (Figures 1–3), we next determined that the addition of J2 fibroblasts results in E2-S23A protein turnover via the proteasome (Figure 4A). Lanes 2 and 3 demonstrate equivalent levels of E2-WT and E2-S23A expression in the absence of J2 cells. Following co-culture with J2 cells, E2-S23A expression is reduced (compare lane 3 with lane 4). Following 6 hours of treatment with the proteasomal inhibitor MG132, E2-S23A levels are detectable in the presence of J2 fibroblasts (lane 7) and this restored expression persists at 8 hours following MG132 treatment (lane 8). This was repeated and quantitated (Figure 4B). Previously, we have demonstrated that the stability of E2 can be regulated by acetylation, and the addition of MG132 restores acetylated E2-S23A (Figure 4C). Acetylated E2 is detected in E2-WT and E2-S23A samples without J2 cells (lanes 3 and 4, respectively), and following 6 and 8 hours of MG132 treatment acetylated levels of E2-S23A are recovered (lanes 8 and 9). We then carried out an IP with E2 and blotted for E2 and acetylated lysine (Figure 4D). This confirmed that when E2 is detectable, it is acetylated. Previously we demonstrated that the class III deacetylase SIRT1 can deacetylate E2 resulting in its destabilization (43). Figure 4E demonstrates that SIRT1 interacts with both E2-WT and E2-S23A (lanes 3 and 4), and that when E2-S23A is stabilized vial the proteasome it is in complex with SIRT1 (lanes 8 and 9). E2-WT protein levels are not affected by MG132 over an 8 hour time period demonstrating that E2-WT is relatively stable in N/Tert-1 cells (Figure 4F). At all times, E2-WT remains acetylated (Figure 4G).

**Figure 4.**
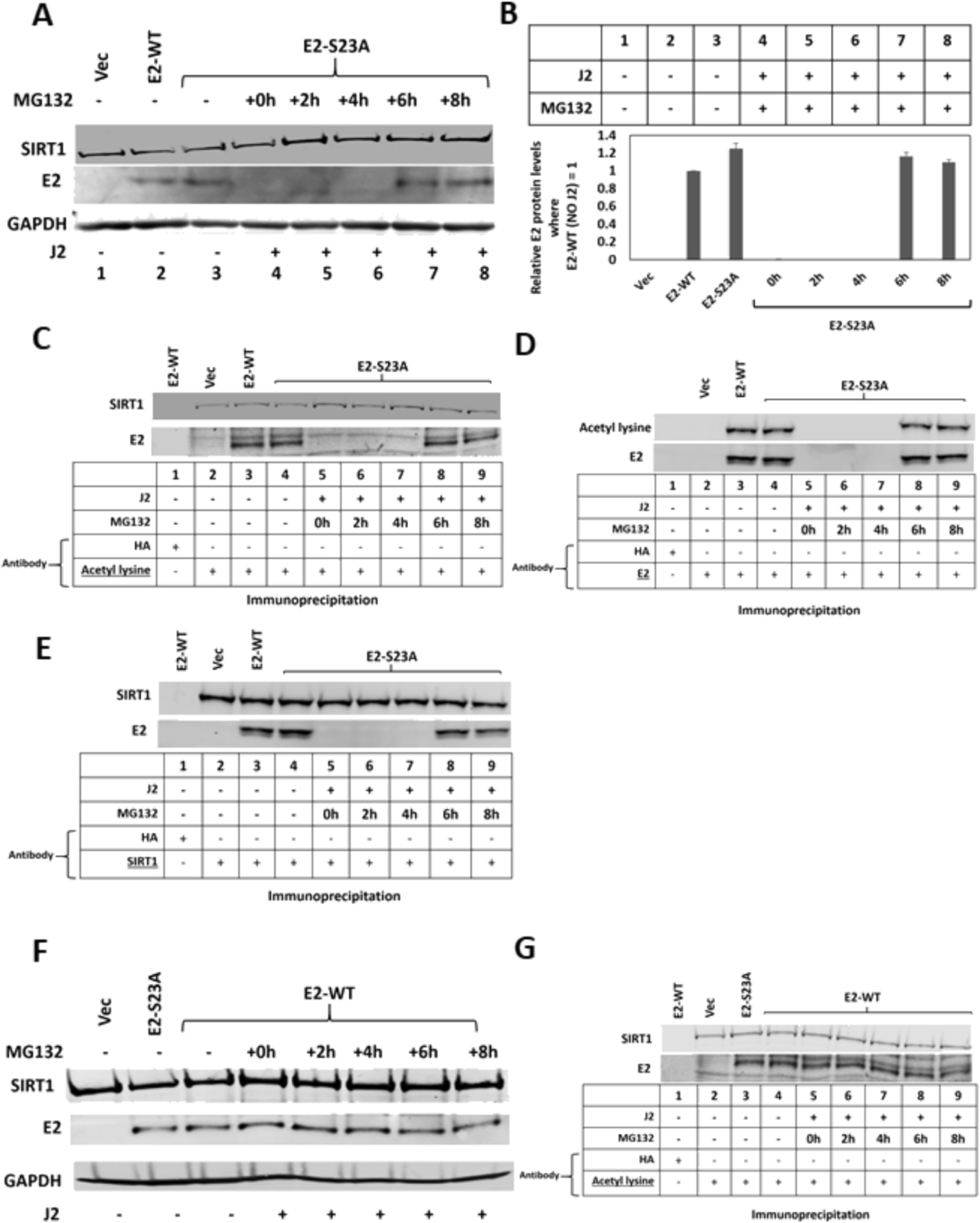
E2-S23A is degraded via the proteasome following the addition of J2 cells. A. Protein extracts were prepared from the indicated cells and western blotted and probed with the indicated antibodies, GAPDH serves as a loading control. The presence of J2 fibroblasts is indicated by the + sign below each lane. The proteasomal inhibitor MG132 was added for the indicated time-period (h = hours) prior to cell harvest and protein preparation. B. The experiment in A was repeated and the levels of E2 proteins in each sample quantitated. C. The protein extracts from the western blot shown in A were immunoprecipitated with an acetylated lysine antibody, demonstrating that all detectable E2 protein is acetylated. D. The protein extracts from the western blot shown in A were immunoprecipitated with a sheep E2 antibody (43) and blotted with the acetyl lysine antibody and a mouse E2 antibody (34). E. The protein extracts from the western blot shown in A were immunoprecipitated with a SIRT1 antibody then blotted for E2 and SIRT1. F. N/Tert-1+E2-WT cells were treated with MG132 for the indicated time points and protein extracts prepared and the indicated western blots carried out. G. E2-WT is acetylated at all time points following MG132 treatment.

Figure 4 suggested that SIRT1 deacetylation of E2-S23A in the presence of J2 cells may be regulating the stability of E2-S23A and we investigated this by knocking down SIRT1 expression. siRNA knockdown of SIRT1 protein levels restores E2-S23A protein levels in the presence of J2 cells (Figure 5A). SIRT1 knockdown has no effect on E2-WT protein levels irrespective of the presence of J2 fibroblasts (lanes 2-5). In the absence of fibroblasts, SIRT1 knockdown does not alter E2-S23A protein levels (compare lane 8 with lane 9). However, in the presence of J2 fibroblasts SIRT1 knockdown restores detectable expression of E2-S23A (compare lane 6 with lane 7). This was repeated and quantitated, demonstrating a significant increase in E2-S23A protein expression in the presence of J2 cells when SIRT1 is knocked down (Figure 5B). The SIRT1 siRNA knockdown and protein quantitation was repeated with an additional SIRT1 siRNA, with identical results (Figure 5C and 5D). We confirmed that the recovery of E2-S23A protein levels in the presence of J2 fibroblasts by SIRT1 knockdown resulted in an acetylated E2-S23A protein (Figure 5E, compare lane 6 with lane 7). Any changes in E2 protein levels detected were not due to alterations in E2 RNA levels with the J2 and SIRT1 knockdown treatments (Figure 5F, results generated using SIRT1 siRNA-A). These results demonstrate that J2 fibroblasts promote the ability of SIRT1 to deacetylate E2-S23A, resulting in proteasomal degradation. Therefore, the E2-TopBP1 interaction is critical for maintaining E2 expression in human keratinocytes interacting with the stroma.

**Figure 5.**
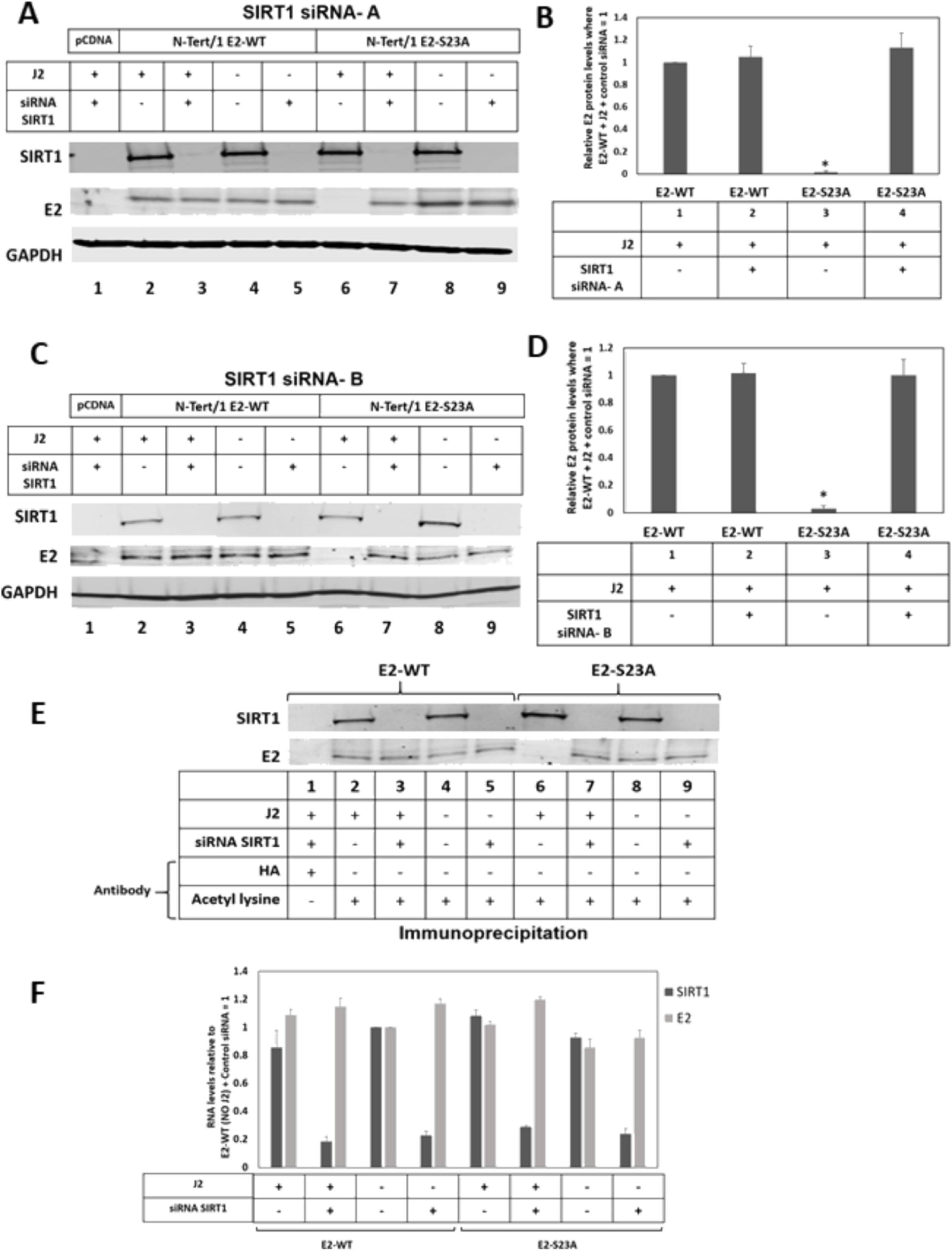
The class III deacetylase SIRT1 is required for the destabilization of E2-S23A in the presence of J2 fibroblasts. A. The indicated cells were treated with scrambled siRNA or siRNA targeting SIRT1 in the presence or absence of J2 fibroblasts. Protein extracts were prepared and western blotting used to monitor SIRT1 and E2 protein levels, GAPDH was used as a loading control. B. The experiment shown in A was repeated and quantitated and the levels of E2-WT protein in the presence or absence of SIRT1with or without J2 fibroblasts calculated. C-D. These figures represent a repetition of the SIRT1siRNA experiments using an alternative SIRT1 targeting siRNA. In B and C, * indicates a statistically significant reduction in E2 protein levels when TopBP1 expression is knocked down in the presence of J2 fibroblasts, p-value < 0.05. E. The protein extracts from the western blot shown in A were immunoprecipitated with an acetylated lysine antibody, demonstrating that knockdown of SIRT1 restores E2-S23A acetylation in the presence of J2 fibroblasts. F. Neither the SIRT1 siRNA treatment nor the addition of J2 fibroblasts statistically altered E2-WT or E2-S23A RNA expression levels. The results shown are a summary from two independent RNA preparations using SIRT1 siRNA A.

### CK2 inhibitor CX4945, which disrupts the E2-TopBP1 interaction, reduces HPV16 and HPV11 E2 protein levels in human keratinocytes

Previously, we demonstrated that siRNA knockdown of CK2, or treatment with the CK2 inhibitor CX4945, disrupts the E2-TopBP1 interaction in N/Tert-1+E2 cells (35). Figure 6A demonstrates that, in the presence of J2 cells, CX4945 reduces E2 protein levels in N/Tert-1+E2. Treatment of N/Tert-1+E2 cells with DMSO (the vector control for CX4945 delivery) did not decrease E2 protein levels (compare lane 2 with lane 4). However, E2 levels were reduced following CX4945 treatment (compare lane 6 with lane 8). This was repeated and quantitated, demonstrating a statistically significant reduction in E2 protein levels in the presence of J2 cells and CX4945 together (Figure 6B).

**Figure 6.**
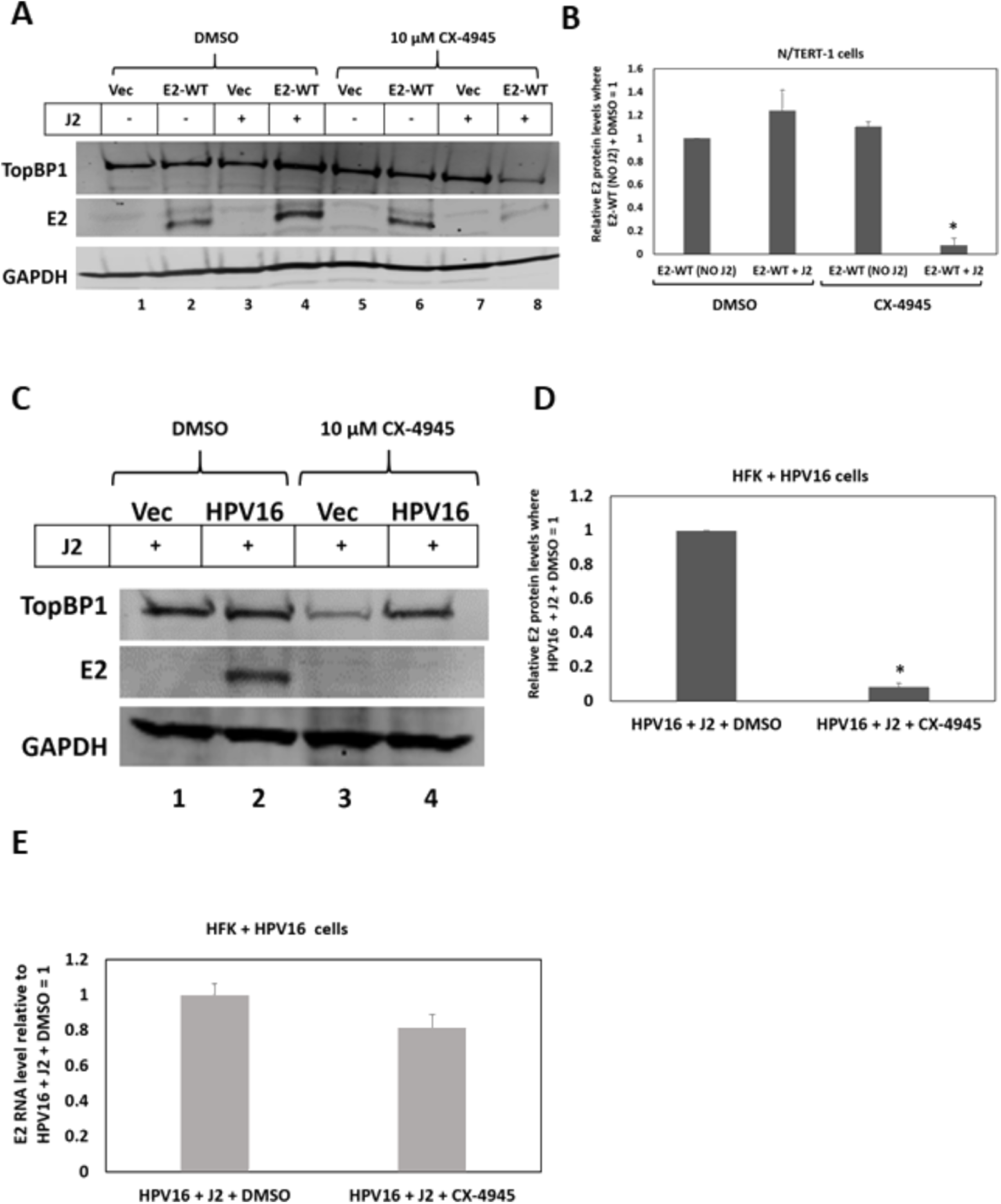
The CK2 inhibitor CX4945, which disrupts the E2-TopBP1 interaction, reduces E2 protein levels in the presence of J2 fibroblasts. A. The indicated cells were treated with DMSO (lanes 1-4) or CK4945 (lanes 5-8) in the presence or absence of J2 fibroblasts. Protein extracts were prepared and western blotting carried out for TopBP1 and E2, GAPDH is used as a loading control. B. The experiment in A was repeated and the levels of E2 protein quantitated. C. The addition of CX4945 reduced E2 protein levels in HFK+HPV16 cells. N/Tert-1+Vec (N/Tert-1 cells selected with G418 using pcDNA3), lanes 1 and 3, was used as a control for E2 expression. In B and D, * indicates a statistically significant reduction in E2 protein levels when CX4945 is added to the HFK+HPV16 cells, p-value < 0.05. E. CX4945 does not significantly alter the levels of E2 RNA expressed in HFK+HPV16 cells; the average of two independent experiments is shown.

Next, we determined that CX4945 can reduce E2 expression in HFK+HPV16 cells in the presence of J2 fibroblasts (Figure 6C). This was repeated and quantitated, demonstrating a significant reduction in E2 protein levels following CX4945 treatment (Figure 6D). We observed a small reduction in E2 RNA levels following CX4945 treatment (Figure 6E), perhaps due to viral genome integration following E2 protein level reduction.

Figures 1–6 demonstrate that interaction with TopBP1 is required for E2 expression during the viral life cycle. When the interaction is disrupted by mutation, knockdown of TopBP1 using siRNA, or CX4945 treatment, E2 is degraded. While CX4945 treatment would be an effective antiviral to prevent infection, disruption of existing infections could promote viral genome integration and therefore progression to cancer. However, for low risk HPV types such as HPV11 or HPV6 that do not cause cancer, targeting E2 degradation could prevent viral genome amplification and therefore reduce viral load. Figure 7A demonstrates stable expression of a tagged HPV11 E2 protein in N/Tert-1 cells. Serine 23, the residue critical for interaction with TopBP1 in HPV16 E2, is conserved in HPV11 E2 and Figure 7B confirms that there is an interaction between HPV11 E2 and TopBP1. In lane 1, immunoprecipitation with an HA (hemagglutinin) epitope antibody, acting as a control, does not pull-down TopBP1 or E2 from N/Tert-1+HPV11 E2 extracts. The TopBP1 antibody pulls down HPV11 E2 (lane 3). siRNA knockdown of TopBP1 reduces the expression of HPV11 E2 in N/Tert-1+HPV11 E2 cells in the presence of J2 cells (Figure 7C). In the absence of J2 cells, knockdown of TopBP1 did not significantly change HPV11 E2 protein levels (compare lane 3 with lane 4). Co-culture with J2 cells did not significantly alter HPV11 E2 levels with control siRNA samples (compare lane 2 with lane 4). However, siRNA knockdown of TopBP1 combined with the addition of J2 fibroblasts reduced E2 protein levels significantly (compare lane 2 with lane 5).

**Figure 7.**
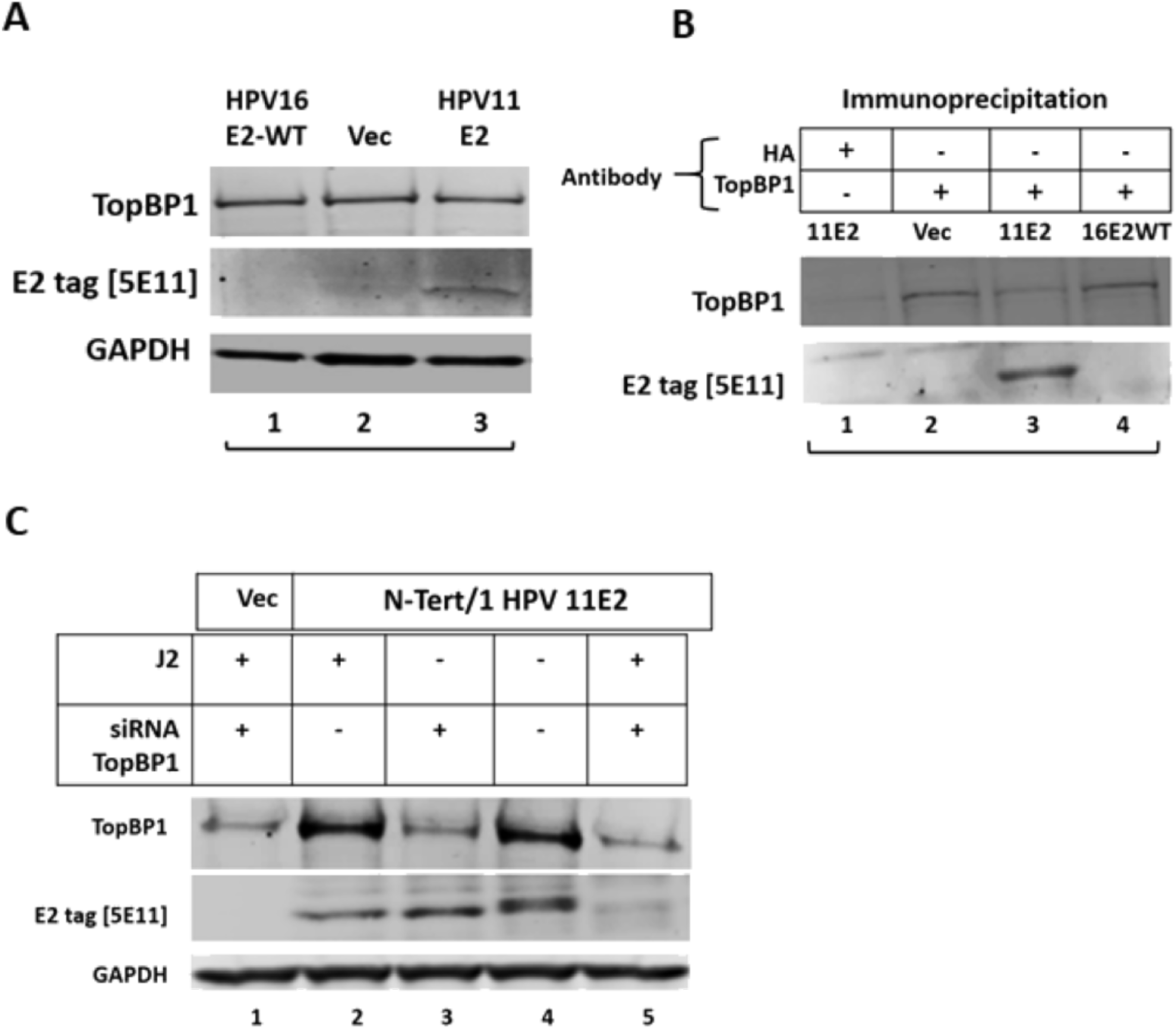
HPV11 E2 protein levels are regulated by TopBP1 in N/Tert-1 cells in a J2 dependent manner. A. N/Tert-1 cells stably expressing HPV11 E2 were generated (lane 3). Vec = N/Tert-1 cells selected with empty pcDNA3 vector. B. HPV11 E2 co-immunoprecipitates with TopBP1. C. siRNA mediated TopBP1 knockdown reduces HPV11 E2 protein levels in a J2 dependent manner.

To determine whether CK2 promotes the HPV11 E2 interaction with TopBP1 we used the CK2 inhibitor CX4945. Figure 8A demonstrates that CX4945 does not change the levels of HPV11 E2 protein in N/Tert-1 cells (compare lane 2 with lane 4). However, CX4945 does disrupt the interaction between HPV11 E2 and TopBP1 (Figure 8B), as it does for HPV16 E2 (35). The combination of CX4945 and the presence of J2 cells resulted in a significant reduction in HPV11 E2 protein levels (Figure 8C). The addition of J2 cells with the treatment vehicle DMSO did not alter the HPV11 E2 protein levels (compare lane 2 with lane 4), but the addition of J2 cells combined with CX4945 resulted in a significant reduction in HPV11 E2 levels (compare lane 6 with lane 8). This was repeated and quantitated to confirm a significant reduction in HPV11 E2 protein levels in the presence of J2 cells plus CX4945 (Figure 8D).

**Figure 8.**
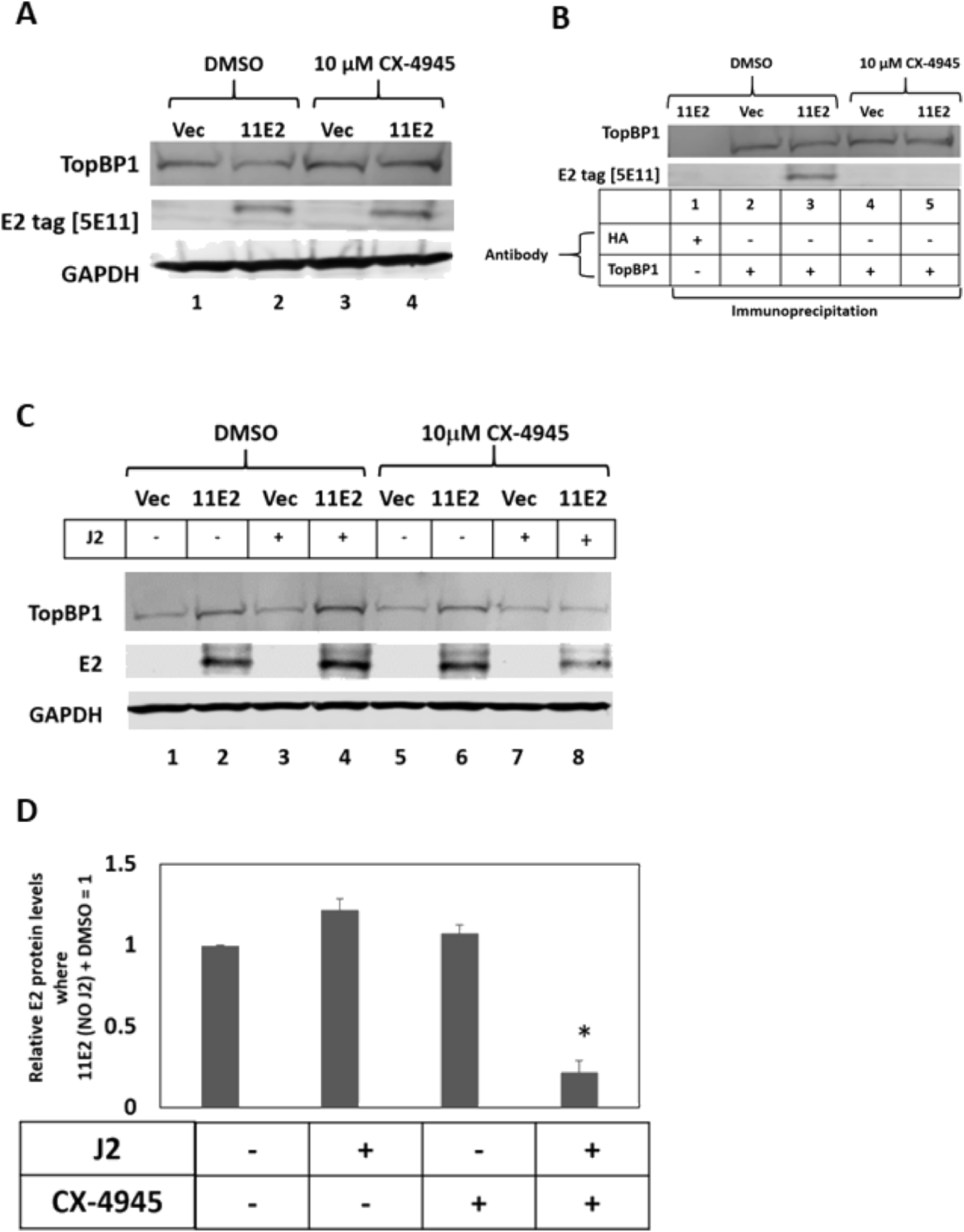
CX4945 reduces HPV11 E2 protein levels in N/Tert-1 cells co-cultured with J2 cells. A. In the absence of J2 fibroblasts, CX4945 does not regulate the expression of HPV11 E2 in N/Tert-1 cells. B. The samples in A were immunoprecipitated with the indicated antibodies and the results demonstrate that CX4945 disrupts the HPV11 E2 interaction with TopBP1 (compare lane 5 with lane 3). C. The indicated cells were treated with DMSO (lanes 1-4) or CK4945 (lanes 5-8) in the presence or absence of J2 fibroblasts. Protein extracts were prepared and western blotting carried out for TopBP1 and HPV11 E2, GAPDH is used as a loading control. D. The experiment in C was repeated and the levels of HPV11 E2 protein quantitated. * indicates a statistically significant reduction in HPV11 E2 protein levels when CX4945 is added to N/Tert-1 cells co-cultured with J2 fibroblasts, p-value < 0.05.

## Discussion

Integration of the HPV16 genome into that of the host provides a cellular growth advantage (44), and tumors with integrated viral DNA have worse clinical outcomes (45–48). Viral genome integration represents the end of the viral life cycle, therefore the virus must remain episomal if it is to propagate. A major controller of the episomal status of the HPV16 genome is the E2 protein, the expression of which is lost during the integration process (45). Not only does E2 regulate transcription from and replication of the viral genome, it is also required for viral genome segregation during mitosis (11). Recently we demonstrated that an interaction between E2 and the cellular protein TopBP1 is critical for the viral genome segregation function of E2 (34, 35). In this model, CK2 phosphorylation of E2 at serine 23 is required for the interaction between E2 and TopBP1. In addition, the interaction between E2 and TopBP1 is required for E2 protein stabilization during mitosis, which is critical for mediating the viral genome segregation function. We introduced an E2 serine 23 mutation to alanine (S23A) in the context of the entire HPV16 genome. During organotypic raft culture, this mutant genome exhibited a more transformed phenotype than the wild type genome (35). There were increased numbers of koilocytes and keratin whorls that are associated with a more transformed cell, and the keratinocytes invaded into the fibroblast-collagen “stroma”, something that never occurred with the wild type HPV16 genomes (35). We saw a lack of E2 positive staining in S23A mutant genome rafts when compared with the wild type, suggesting that there has been a loss of E2 protein during the rafting process. The loss of E2 expression would abolish viral replication and promote viral genome integration, explaining the more transformed phenotypes observed with the S23A organotypic raft samples.

In this report we demonstrate that, in human foreskin keratinocytes immortalized by the S23A genome (HFK+HPV16-E2-S23A), there was a lack of detectable E2 protein expression due to instability that resulted in viral genome integration during organotypic raft culture. Using N/Tert-1 lines stably expressing E2 and E2-S23A, we demonstrate that the addition of J2 fibroblasts promotes E2-S23A degradation via the proteasome and that SIRT1 knockdown can rescue E2-S23A expression. SIRT1 is a class III deacetylase that we have already demonstrated deacetylates E2 and regulates its stability (43). To our knowledge, this is the first description of stromal regulation of SIRT1 function. Figure 9 summarizes our proposed model. E2 interaction with TopBP1 controls the ability of SIRT1 to deacetylate E2 (upper panel); SIRT1 is an interacting partner of TopBP1 and can regulate its acetylation status (49, 50). However, in the absence of TopBP1 interaction, J2 fibroblasts hyper-activate SIRT1 resulting in complete deacetylation (lower panel). Alternatively, the fibroblasts could attenuate the function of an acetylase that would allow SIRT1 to increase the deacetylation of E2. We propose that this promotes ubiquitination of a lysine residue (perhaps the same one that is acetylated) resulting in proteasomal degradation. Therefore, when not complexed with TopBP1, E2-S23A is vulnerable to stromal mediated degradation. In the HPV31 life cycle, the knockdown of SIRT1 using shRNA results in HPV31 viral genome integration, perhaps because there has been an alteration in the HPV31 E2 protein levels; this requires further investigation (51). Another critical role for SIRT1 is the regulation of WRN acetylation, a process that is important in maintaining the fidelity of E1-E2 DNA replication (52). Overall, these results demonstrate that E2 stability is controlled via host interactions, and that the interaction with TopBP1 is critical for E2 expression and therefore for the viral life cycle.

**Figure 9.**
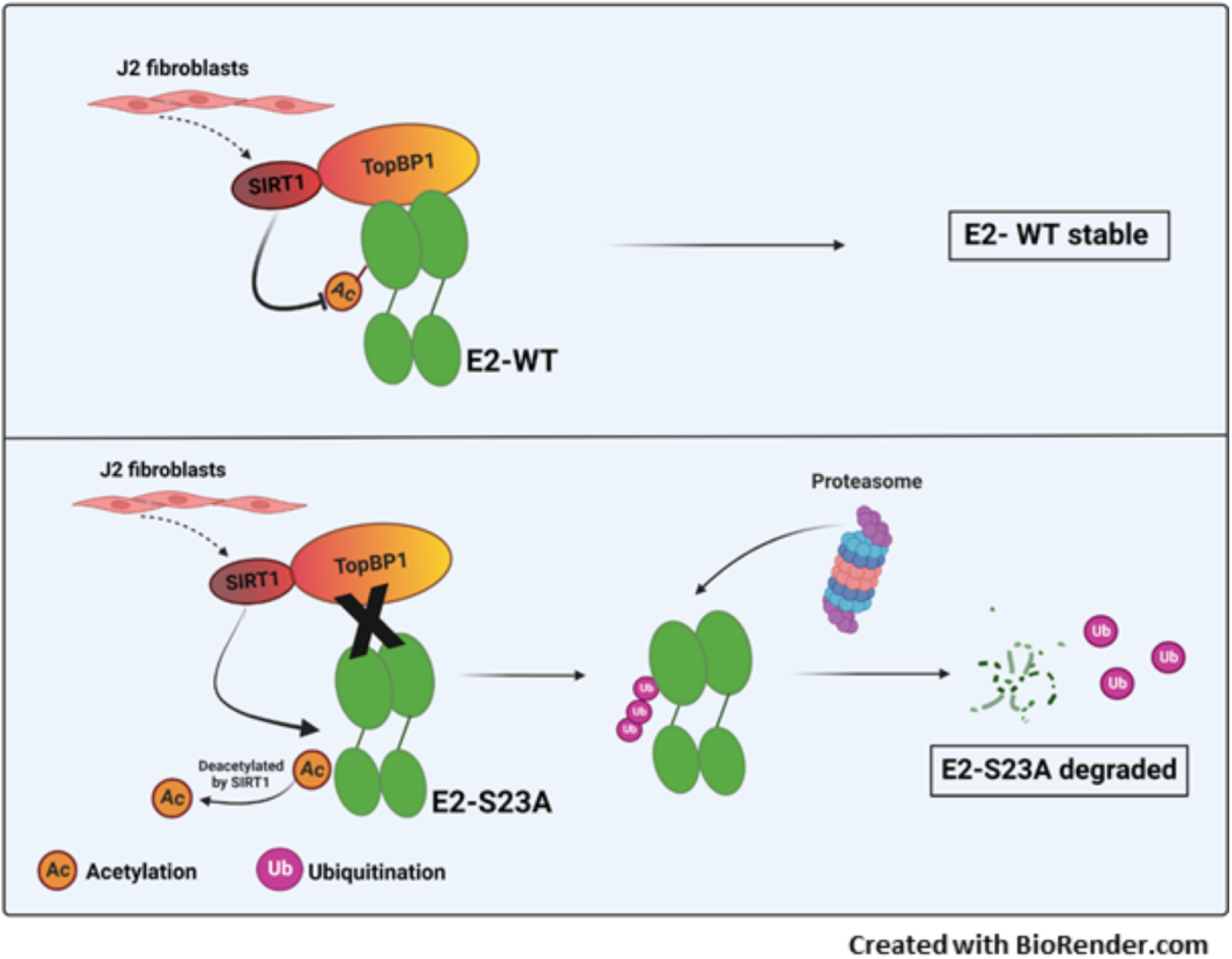
A model explaining the regulation of E2 protein stability in the presence of J2 fibroblasts. In the top panel E2 is complexed with TopBP1 and the ability of SIRT1 to deacetylate E2 is regulated, allowing a stable level of E2 expression. In the lower panel, if the E2 interaction with TopBP1 is disrupted (by E2 mutation, siRNA treatment of TopBP1, or the addition of CX4945 to disrupt the E2-TopBP1 interaction) then SIRT1 deacetylation of E2 is accelerated. This deacetylation promotes ubiquitination and subsequent proteasomal degradation of the E2 protein.

Recently, we demonstrated that TopBP1 and BRD4 are in the same cellular complex, and it is likely that both of these proteins are involved in mediating the plasmid segregation function of E2 (currently under investigation). Another striking feature of the E2 interaction with BRD4 is that it can also regulate E2 protein stability (53, 54). Overexpression of the BRD4 carboxyl-terminal domain (CTD) results in E2 protein stabilization and disruption of HPV16 E2 interaction with the cullin-3 complex (54). Recently, we demonstrated E2 stabilization during mitosis (irrespective of J2 fibroblasts), and that this stabilization is required for the plasmid segregation function of E2 (34, 35).

CK2 has been proposed as a cancer therapeutic target, and CX4945 has been in clinical trial (55). Treatment with this drug has the potential to destabilize HPV16 E2 as it would disrupt interaction with TopBP1 and promote integration of the HPV16 genome, which could promote HPV16 cancer development. However, for low risk HPV, such as HPV11, CK2 inhibitors potentially offer a therapeutic window. Here we demonstrate that CX4945 treatment of human keratinocytes disrupts HPV11 E2 interaction with TopBP1, and destabilizes HPV11 E2 in the presence of J2 fibroblasts. Serine 23 on HPV16 E2 that we demonstrated is phosphorylated by CK2 is conserved in both HPV11 and HPV6 E2 proteins. Juvenile respiratory papillomatosis is caused primarily by HPV11 and HPV6 infections and the standard of care for this disease remains removal of the papillomas (56). The degradation of HPV11 E2 by CX4945 represents a therapeutic opportunity for the management of this disease. The reduction of E2 would disrupt viral genome replication and therefore viral production, potentially alleviating the disease burden in patients suffering from juvenile respiratory papillomatosis. Inhibition of CK2 by CX4945 also regulates the stability and nuclear retention of HPV11 and HPV18 E1 adding another potential mechanism for CX4945 targeting of HPV replication (57).

We are currently investigating the mechanisms that control HPV16 E2 stability throughout the cell cycle, and the host factors that mediate this stabilization. Disruption in expression of, or mutation of, these host factors could disrupt E2 stability and promote viral genome integration. Therefore, the complex that regulates E2 stability is critical for the viral life cycle, and disruption of this complex could promote viral integration and HPV16 cancer progression. Overall, the results demonstrate the crucial nature of the E2-TopBP1 interaction during the viral life cycle and in HPV16 disease progression.

## Materials and methods

### Cell culture, plasmids, and reagents

Stably expressing HPV16 wild type E2 (E2-WT), E2-S23A (E2 with serine 23 mutated to alanine, abrogating interaction with TopBP1) and pcDNA empty vector plasmid control were generated in N/Tert-1 cells as described previously (34, 35). Stable HPV11 E2 cell lines was also generated in N/Tert-1 cells using the HPV11 E2 expression vector, which was a kind gift from Dr. Reet Kurg (58). N/Tert-1 cells were cultured in keratinocyte serum-free medium (K-SFM) (Invitrogen; catalog no. 37010022) supplemented with bovine pituitary extract, EGF (Invitrogen), 0.3 mM calcium chloride (Sigma; 21115) and 150 μg/ml G418 (Thermo Fisher Scientific) at 37 °C in a 5% CO_2_/95% air atmosphere. Human foreskin keratinocytes (HFKs) were immortalized with HPV16 (WT and S23A) and cultured as previously described (35). Mitomycin C-treated 3T3-J2 fibroblasts feeders were plated 24 hours prior to plating N/Tert-1 or HFK cells on top of the feeders, in their respective cell culture media and allowed to grow to 70% confluency.

### Organotypic raft culture and protein isolation from rafts

Keratinoctye lines containing wild-type or mutant HPV16 genomes, encoding the E2-S23A mutation, were differentiated via organotypic raft culture as described previously (ref). Briefly, cells were seeded onto type 1 collagen matrices containing J2-3T3 fibroblast feeder cells. Keratinocytes were then grown to confluency atop the collagen matrices, which were then lifted onto wire grids and cultured in cell culture dishes at the air-liquid interface, with media replacement on alternate days. Following 7 or 14 days of culture, cultures were peeled off of collagen matrix and homogenized using a Dounce homogeniser in either HIRT buffer (0.6% SDS, 10 mM EDTA, 50 mM NaCl pH 7.5), or urea lysis buffer (8 M urea, 50 mM Tris-HCl pH 8, 0.15 M β-mercaptoethanol, supplemented with protease and phosphatase inhibitors). Following homogenization, DNA samples were treated with RNase A before digestion with proteinase K. DNA was then extracted using phenol chloroform, as previously described (59). For protein, homogenized samples were incubated on ice for an hour with frequent agitation. The soluble and insoluble fractions were separated by centrifugation before Western blot analysis.

### Western blotting

Specified cells were trypsinized, washed with 1X PBS and resuspended in 2x pellet volume protein lysis buffer (0.5% Nonidet P-40, 50 mM Tris [pH 7.8], 150 mM NaCl) supplemented with protease inhibitor (Roche Molecular Biochemicals) and phosphatase inhibitor cocktail (Sigma). Cell pellet-buffer suspension was incubated on ice for 20 min and afterwards centrifuged for 20 min at 184,000 rcf at 4 °C. Protein concentration was determined using the Bio-Rad protein estimation assay according to manufacturer’s instructions. 50 μg protein was mixed with 4x Laemmli sample buffer (Bio-Rad) and heated at 95 °C for 5 min. Protein samples were separated on Novex 4–12% Tris-glycine gel (Invitrogen) and transferred onto a nitrocellulose membrane (Bio-Rad) at 30V overnight using the wet-blot transfer method. Membranes were then blocked with Odyssey (PBS) blocking buffer (diluted 1:1 with 1X PBS) at room temperature for 1-hour and probed with indicated primary antibody diluted in Odyssey blocking buffer, overnight. Membranes were washed with PBS supplemented with 0.1% Tween (PBS-Tween) and probed with the Odyssey secondary antibody (goat anti-mouse IRdye 800CW or goat anti-rabbit IRdye 680CW) (Licor) diluted in Odyssey blocking buffer at 1:10,000. Membranes were washed twice with PBS-Tween and an additional wash with 1X PBS. After the washes, the membrane was imaged using the Odyssey® CLx Imaging System and ImageJ was used for quantification, utilizing GAPDH as internal loading control. Primary antibodies used for western blotting studies are as follows: monoclonal B9 1:500 (60), TopBP1 1:1,000 (Bethyl, catalog no. A300-111A), GAPDH 1:10,000 (Santa Cruz; catalog no. sc-47724), SIRT1 antibody 1:1000 (Sigma, catalog no. 07–131), phospho-γH2AX 1:1000 (Cell Signaling Technology, catalog no. 9718), HPV16 E7 1:200 (Santa Cruz, catalog no. sc-6981), E2 tag [5E11] 1:1000 (Abcam, catalog no. ab977).

### Immunoprecipitation assay

Cell lysate was prepared as described above. 250 μg of the lysate was incubated with lysis buffer (0.5% Nonidet P-40, 50mM Tris [pH 7.8], and 150mM NaCl), supplemented with protease inhibitor (Roche Molecular Biochemicals) and phosphatase inhibitor cocktail (Sigma) to a total volume of 500 μl. Primary antibody of interest or a HA-tag antibody (used as a negative control) was added to this prepared lysate and rotated at 4°C overnight. The following day, 40 μl of protein A beads per sample (Sigma; equilibrated to lysis buffer as mentioned in the manufacturer’s protocol) was added to the above mixture and rotated for another 4 hours at 4°C. The samples were gently washed with 500 μl lysis buffer by centrifugation at 1000 x g for 2-3 min. This wash was repeated three times. The bead pellet was resuspended in 4X Laemmli sample buffer (Bio-Rad), heat denatured and centrifuged at 1000 x g for 2-3 min. The supernatant was gel electrophoresed using an SDS-PAGE system which was later transferred onto a nitrocellulose membrane using wet-blot transfer method. The membrane was probed for the presence of E2, SIRT1 or TopBP1, as mentioned in the description of western blotting above.

### Southern blotting

Total cellular DNA was extracted using a phenol chloroform method and 5 micrograms digested with either *SphI* or *HindIII*, to linearise the HPV16 genome or leave episomes intact, respectively. All digests included *DpnI* to ensure that all input DNA was digested and not represented as replicating viral DNA. Digested DNA was separated by electrophoresis of a 0.8% agarose gel, transferred to a nitrocellulose membrane and probed with radiolabeled (32-P) HPV16 genome. This was then visualized by exposure to film.

### RNA isolation and SYBR green qRT-PCR

The SV total RNA isolation system kit (Promega) was employed to isolate RNA from cells, as per the manufacturer’ protocol. A high-capacity cDNA reverse-transcription kit from Invitrogen was used to synthesize cDNA, which was processed for qPCR.

### Real-time PCR (qPCR)

qPCR was performed on 10 ng of DNA Hirt extracted from organotypic raft grown cells or on the cDNA isolated, as described above. DNA and relevant primers were mixed with PowerUp SYBR green master mix (Applied Biosystems), and real-time PCR was performed using the 7500 Fast real-time PCR system, using SYBR green reagent. Expression was quantified as relative quantity over GAPDH using the 2^-ΔΔCT^ method. Primer used are as follows. HPV16 E2 F, 5’-ATGGAGACTCTTTGCCAACG-3’; HPV16 E2 R, 5’ TCATATAGACATAAATCCAG-3’; HPV16 E6 F, 5’-TTGAACCGAAACCGGTTAGT-3’; HPV16 E6 R,5’-GCATAAATCCCGAAAAGCAA-3’; HPV16 E7 F, 5’ - ATATATGTTAGATTTGCAACCAGAGACAAC −3’; HPV16 E7 R, 5’ −GTCTACGTGTGTGCTTTGTACGCAC −3’; TopBP1 F, 5’-TGAGTGTGCCAAGAGATGGAA-3’; TopBP1 R, 5’-TGCTTCTGGTCTAGGTTCTGT-3’; SIRT1 F, 5’-CAGTGTCATGGTTCCTTTGC-3’; SIRT1 R, 5’-CACCGAGGAACTACCTGAT-3’; Glyceraldehyde-3-phosphate dehydrogenase (GAPDH) F, 5’-GGAGCGAGATCCCTCCAAAAT-3’ and GAPDH R, 5’-GGCTGTTGTCATACTTCTCATGG-3’.

### MG132 proteasomal inhibitor treatment

N/Tert-1 cells expressing stable E2-S23A were plated on mitomycin C-treated J2 feeders in KSFM media. Next day, the cells were treated with 10μM of MG132 (Z-Leu-Leu-Leu-al; Sigma, catalog no. C2211) and harvested at 0, 2, 4, 6, and 8 hours post-treatment and immunoblotting was carried out to detect E2, SIRT1 and GAPDH.

### CK2 inhibitor treatment

N/Tert-1 and HFK cells were plated at a density of 3 x 10^5^ in a 100-mm plate. Following day, the cells were treated with 10 μM CK2 inhibitor, CX-4945 (Silmitasertib; APExBIO, catalog no. A8330) or 10 μM DMSO for 48 hours. The cells were then harvested and processed for western blotting to detect E2, TopBP1 and GAPDH as described in the section above.

### siRNA treatment

N/Tert-1 cells were plated on a 100-mm plates. The next day, cells were transfected with 10 μM of the siRNA mentioned below. 10 μM of MISSION® siRNA Universal Negative Control (Sigma-Aldrich; catalog no. SIC001) was used as a “non-targeting” control in our experiments. Lipofectamine™ RNAiMAX transfection (Invitrogen, catalog no. 13778-100) protocol was used in the siRNA knockdown. 48 hours post transfection, the cells were harvested, and the knockdown was confirmed by immunoblotting for the protein of interest and cells were also processed for qRT-PCR, as described above. All siRNAs were purchased from Sigma-Aldrich: siRNA TopBP1-A-CUCACCUUAUUGCAGGAGA; siRNA TopBP1-B-GUAAAUAUCUGAAGCUGUA; siRNA SIRT1-A-ACUUUGCUGUAACCCUGUA; siRNA SIRT1-B-AGAGUUGCCACCCACACCU.

### Statistical analysis

Results are represented as means ± SE. Standard error was calculated from three independent experiments. Student’s *t* test was used to determine significance.

## Acknowledgements

This work was supported by VCU Philips Institute for Oral Health Research, National Cancer Institute designated Massey Cancer Center grant P30 CA016059, and R01 DE029471 (IMM).

